# Chloroquine inhibits Zika Virus infection in different cellular models

**DOI:** 10.1101/051268

**Authors:** Rodrigo Delvecchio, Luiza M Higa, Paula Pezzuto, Ana Luiza Valadão, Patrícia P Garcez, Fábio L Monteiro, Erick C. Loiola, Stevens Rehen, Loraine Campanati, Renato Santana de Aguiar, Amilcar Tanuri

## Abstract

Zika virus (ZIKV) infection *in utero* might lead to microcephaly and other congenital defects. In adults, cases of Guillain-Barré syndrome and meningoencephalitis associated with ZIKV infection have been reported, and no specific therapy is available so far. There is urgency for the discovery of antiviral agents capable of inhibiting viral replication and its deleterious effects. Chloroquine is widely administered as an antimalarial drug, anti-inflammatory agent, and it also shows antiviral activity against several viruses. Here we show that chloroquine exhibits antiviral activity against ZIKV in VERO, human brain microvascular endothelial, and neural stem cells. We demonstrated *in vitro* that chloroquine reduces the number of ZIKV-infected cells, virus production and cell death promoted by ZIKV infection without cytotoxic effects. Our results suggest that chloroquine is a promising candidate for ZIKV clinical trials, since it is already approved for clinical use and can be safely administered to pregnant woman.

## INTRODUCTION

Zika virus (ZIKV) is an arthropod-borne virus, transmitted by *Aedes* mosquitoes, that belongs to the *Flavivirius* genus, which also includes other pathogens such as West Nile Virus (WNV), Yellow Fever Virus (YFV), Japanese Encephalitis Virus (JEV) and Dengue Virus (DENV). Phylogenetic analysis of the nonstructural protein 5 encoding region identified three ZIKV lineages: East African, West African and Asian (Lanciotti et al., 2008).

*In utero* exposure to ZIKV might lead to microcephaly and other developmental malformations including calcifications, arthrogryposis, ventriculomegaly, lissencephaly, cerebellar atrophy and ocular abnormalities, which altogether are referred as congenital Zika syndrome (Brasil et al., 2016; Martines et al., 2016; Mlakar et al., 2016; Oliveira Melo et al., 2016). Although all ZIKV lineages can infect humans, severe manifestations of the infection have only been associated to Asian lineages, including Brazilian isolates (Calvet et al., 2016; Mlakar et al., 2016).

An unprecedented increase in microcephaly cases associated with ZIKV infection prompted the World Health Organization (WHO) to declare it a Public Health Emergency of International Concern. Brazil is currently the most affected country with 1,168 cases confirmed of ZIKV-related microcephaly (http://portalsaude.saude.gov.br).

ZIKV was detected in the brain and amniotic fluid of newborns and stillborns with microcephaly (Brasil et al., 2016; Calvet et al., 2016; Martines et al., 2016; Mlakar et al., 2016) and it was shown to kill human neuroprogenitor cells *in vitro* as well as decrease the brain organoid growth rate (Garcez et al., 2016; Tang et al., 2016). After reviewing several evidences, the Centers for Disease Control and Prevention (CDC) concluded that Zika virus infection causes microcephaly and other congenital defects (Rasmussen et al., 2016).

Symptoms of ZIKV infections include low-grade fever, headache, rash, conjunctivitis, arthritis and myalgia (Brasil et al., 2016; Duffy et al., 2009). However, in a minor number of cases, infection is associated with cases of Guillain-Barré syndrome (Cao-Lormeau et al., 2016) and meningoencephalitis (Carteaux et al., 2016). Currently, there is no vaccine or specific therapeutic approaches to prevent or treat ZIKV infections. With the alarming increase in the number of countries affected and the potential for viral spread through global travel and sexual transmission (D’Ortenzio et al., 2016; Deckard et al., 2016), there is an urgency to find a treatment capable of lessen the effects of the disease and inhibit further transmission.

Chloroquine, a 4-aminoquinoline, is a weak base that is rapidly imported into acidic vesicles increasing their pH (Browning, 2014). It is approved by the Food and Drug Administration (FDA) to treat malaria and has long been prophylactically prescribed to pregnant women at risk of exposure (Levy et al., 1991). Chloroquine, through inhibition of pH-dependent steps of viral replication, restricts HIV (Tsai et al., 1990), Influenza virus (Ooi et al., 2006), DENV (Juvenal et al., 2013), JEV (Zhu et al., 2012) and WNV infection (Boonyasuppayakorn et al., 2014). Here, we sought to investigate the antiviral effects of chloroquine on ZIKV infection in different cell types.

## RESULTS

### ZIKV infection is inhibited by chloroquine in Vero cells

We have initially characterized the antiviral properties of chloroquine in Vero cells, a model widely used to study viral infections. Vero cells were infected with ZIKV MR766 at MOI 2 and then treated with chloroquine in concentrations ranging from 6.25 to 50 μM for 5 days. Viral infectivity was assessed using 4G2 antibody, which detects flavivirus envelope protein. We observed that chloroquine treatment decreases the number of ZIKV-infected cells in a dose dependent manner. Flow cytometry analysis showed a reduction of 65% and 95% in ZIKV-infected cells treated with 25 μM and 50 μM chloroquine, respectively, compared to untreated infected cells (Fig. 1A). These results were corroborated by immunofluorescence staining (Fig. 1B). Additionally, chloroquine decreases the production of infectious virus particles by ZIKV-infected cells (Fig. 1C). To confirm that viral inhibition is independent of chloroquine cytotoxicity, cell viability was analyzed in uninfected cells treated with chloroquine (1.56 to 200 μM) for 5 days. Chloroquine does not impact cell viability at 50 μM or lower concentrations (Fig. 1D). We further analyzed whether chloroquine treatment could protect Vero cells from ZIKV-induced cell death. Chloroquine, ranging from 12.5 to 50 μM, rescued cell viability to 55-100% (Fig. 1E).

**Figure 1.**
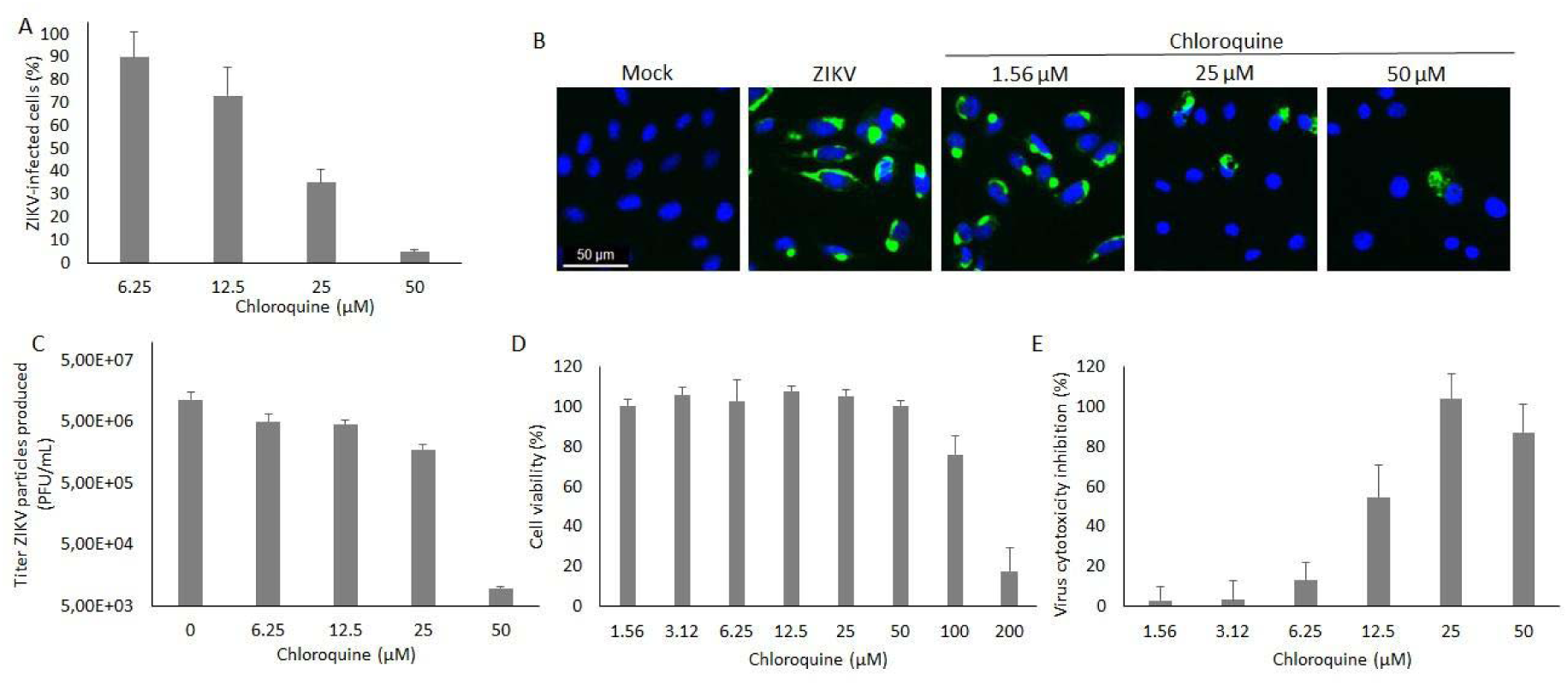
Inhibition of ZIKV infection by chloroquine in Vero cells. Vero cells were infected with ZIKV MR766 at MOI 2 and treated with chloroquine for 5 days when cells were stained for the viral envelope protein and analyzed by flow cytometry (A). (B) Chloroquine treatment of ZIKV-infected cells was evaluated by immunofluorescence staining of 4G2 antibody (green) and DAPI (blue). (C) Infectious virus particles were quantified on supernatant. (D) Chloroquine cytotoxicity was evaluated by cell viability of uninfected cells treated with chloroquine. (E) Protection against ZIKV-induced cell death was evaluated in ZIKV-infected Vero cells treated with chloroquine for 5 days. Data were normalized by each experiment control. Data are represented as mean ± SD.

### Chloroquine reduces ZIKV infection in hBMEC, an *in vitro* model of blood-brain barrier

Considering that ZIKV infects human brain microvascular endothelial cells (hBMECs) (Bayer et al., 2016), we investigated whether chloroquine could inhibit viral infection of these cells. Chloroquine reduces the number of ZIKV-infected hBMECs to 20% and 8% at 25 and 50 μM, respectively (Fig. 2A and D). These concentrations are non-cytotoxic (Fig. 2B) and protected around 80% of hBMECs from ZIKV-induced death (Fig. 2C).

**Figure 2.**
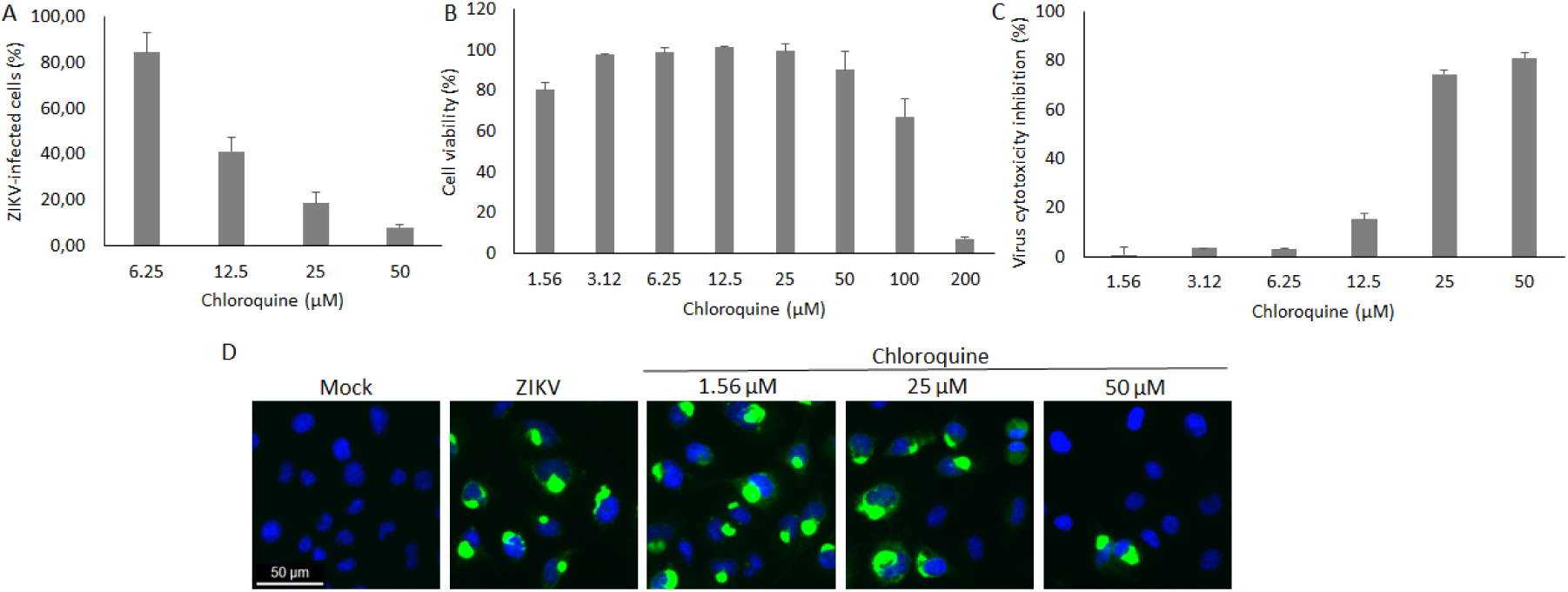
Chloroquine reduces the number of ZIKV-infected hBMEC and protects hBMEC from ZIKV-induced cell death. (A) hBMEC were infected with ZIKV MR 766 at MOI 2 followed by chloroquine treatment for 5 days. Cells were stained with 4G2 antibody and analyzed by flow cytometry. (B) Uninfected hBMECs were incubated with chloroquine for 5 days and cell viability was analyzed. (C) Protection against ZIKV-induced cell death was measured in chloroquine-treated, ZIKV-infected cells. (D) Immunofluorescence with 4G2 antibody (green) and DAPI (blue) of ZIKV-infected cells treated with chloroquine for 5 days. Data are represented as mean ± SD.

### Chloroquine inhibits ZIKV infection in human neural progenitor cells

Neural stem cells (NSCs) are key cells in the process of corticogenesis, giving rise to the three main cell types of central nervous system: neurons, astrocytes and oligodendrocytes. Primary microcephaly occurs mainly as a result of the depletion of the NSCs pool (Gilmore and Walsh, 2013). In order to evaluate if chloroquine could protect these cells from ZIKV infection, they were exposed to up to 50μM chloroquine for 4 days. Chloroquine treatment decreases 60% of the number of ZIKV-infected hNSC cells and protects 70% of these cells from ZIKV cytopathic effects, reducing death without cytotoxicity (Fig. 3A-D).

**Figure 3.**
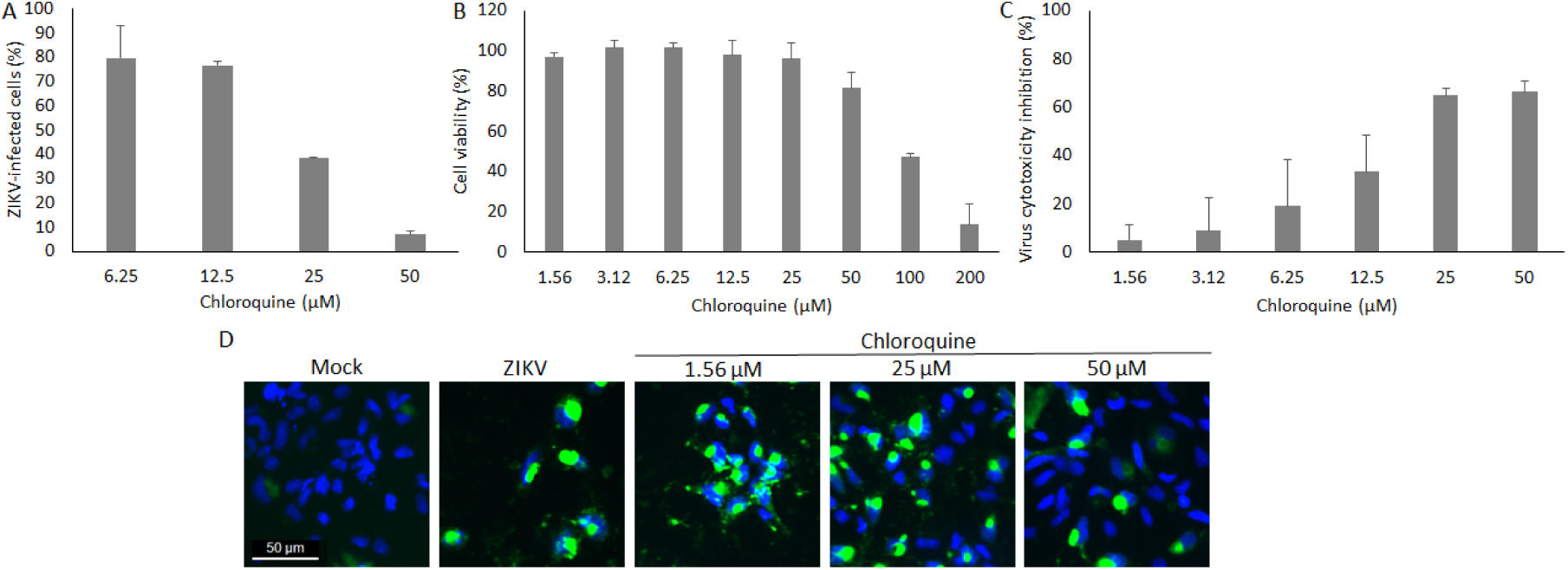
Chloroquine inhibits ZIKV infection in human NSCs. NSCs were infected with ZIKV MR 766 at MOI 2 and incubated with increasing concentrations of chloroquine for 5 days. (A) The number of ZIKV-infected cells was analyzed by 4G2 staining and flow cytometry. (B) Chloroquine cytotoxicity was assessed by the viability of uninfected NSC treated with chloroquine. (C) Chloroquine treatment protection from deleterious effects of infection was evaluated after 5 days of infection. (D) Immunofluorescence of infected and treated cells with 4G2 antibody (green) and DAPI (blue). Data are represented as mean ± SD.

### Chloroquine inhibits ZIKV infection in mouse neurospheres

Neuroprogenitor cells when submitted to differentiation culture conditions generates neurospheres and induced to differentiate into neurons. Our group showed that ZIKV infection affects neurospheres size, neurite extension and neuronal differentiation (Campanati et al., 2016). As we previously observed, neurospheres infected with MR766 ZIKV showed convoluted and misshappen neurites. Neurite extension was evaluated in chloroquine treated cultures by Map2 staining and phase contrast and although many neurospheres were severely impacted by the infection, many other showed the same aspect of mock-infected spheres indicating that choroquine treatment rescued neurite extension phenotype (Fig. 4A-C). ZIKV infection decreased when 12.5 μM chloroquine was added to the medium, as evaluated by 4G2 staining (Fig. 4D-F).

**Figure 4.**
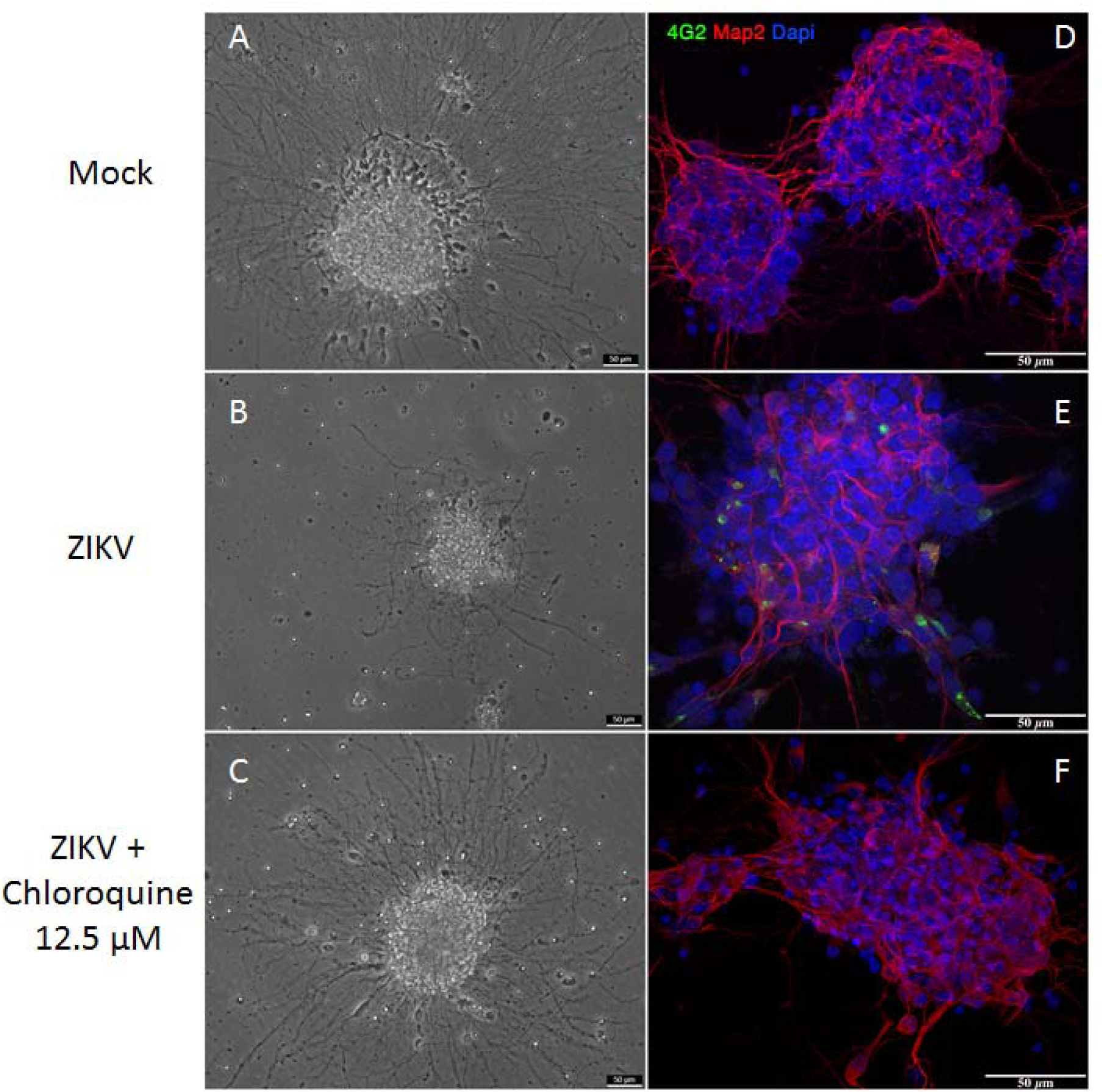
Chloroquine inhibits ZIKV infection in mouse neurospheres. Mouse neurospheres were infected with ZIKV MR766 and treated with chloroquine for 3 days. Neurospheres were analyzed by phase contrast microscopy (A-C) and triple stained for envelope viral protein (green), Map-2 (red), a neuron-specific protein, and DAPI (blue) (D-F).

### Chloroquine inhibits Asian and African ZIKV strains infection

Microcephaly cases and neurological disorders have only been associated to the Asian strains of ZIKV, detected in the French Polynesia and in the Americas (Calvet et al., 2016; Cao-Lormeau, 2014; Mlakar et al., 2016). To determine the inhibition spectrum of chloroquine against ZIKV infection, Vero cells were infected with the African lineage MR766 and two Brazillian isolates of the Asian lineage (ZIKV BR1 and ZIKV BR2). The levels of viral RNA in the supernatant of Vero cells were determined as a direct measurement of ZIKV infection. Treatment with 25 or 50 μM chloroquine led to a 30 to 40-fold reduction in ZIKV particle production, regardless the viral lineage used (FIG. S1).

### Chloroquine inhibits early stages of ZIKV infection

Inhibition of viral infection mediated by chloroquine can occur in both early and later stages of ZIKV replication cycle (Savarino et al., 2003). To evaluate which step of the viral cycle was susceptible to inhibition, chloroquine was added to Vero cells at different time points post-infection. Supernatant was collected 30 hours post-infection and virus production was evaluated by relative quantification of viral RNA over untreated control by qPCR. Virus titers were also determined by plaque assay in Vero cells. Incubation of Vero cells with chloroquine at 0 hpi had a greater impact on production of ZIKV particles, decreasing 70 times viral RNA over control. Addition of chloroquine from 30 minutes to 12 hours post-infection was able to reduce virus release 9-20 times over untreated, infected-cells. However, chloroquine added at 24 hpi had no effect on viral production (FIG. 5A). These results were confirmed by quantification of ZIKV infectious particles release after chloroquine treatment (FIG. 5B). These data confirm that chloroquine targets mainly early stages of viral infection.

**Figure 5.**
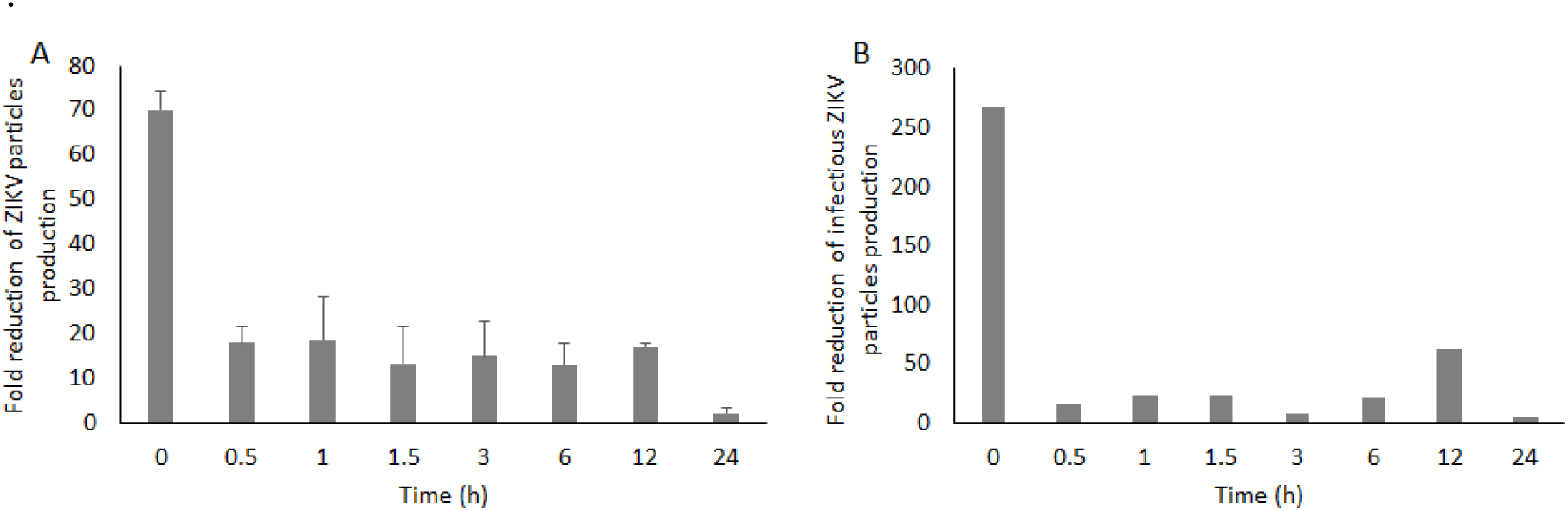
Early stages of infection are inhibited by chloroquine. Vero cells were infected with ZIKV MR766 at MOI 10 and chloroquine added at different times post-infection when the supernatant was collected and viral RNA (A) or infectious particles (B) were quantified.

## DISCUSSION

From January 2007 to April 2016, ZIKV transmission has been reported in 64 countries and territories (WHO, 2016). Although Zika virus disease is in general mild, the recent correlation between infection and congenital malformations and neurological damages in adults has intensified the need for therapeutical approaches. Prophylactic treatment for women that intend to get pregnant in epidemic areas and travelers going to affected countries would represent relevant tools to reduce ZIKV transmission and avoid the spread of the disease by travelers. Moreover, a drug that blocks placental transfer of the virus could decrease the chance of vertical transmission in viremic pregnant women as was shown for HIV-infected pregnant women treated with anti-retroviral therapy (Connor et al., 1994).

Here we demonstrate that chloroquine decreases the number of ZIKV-infected cells and protected them from ZIKV-induced cell death at non-cytotoxic concentrations. The half maximal effective concentration (EC50) of chloroquine, concentration that protected 50% of cells from ZIKV-induced death, was 9.82-14.2 μM depending on cell model and 50% cytotoxicity concentration (CC50) ranged from 94.95-134.54 μM (Table 1). These values of EC50 are lower than those obtained for DENV inhibition (around 25 μM) and HIV inhibition (100 μM) (Juvenal et al., 2013; Tsai et al., 1990).

**Table 1.** Pharmacological parameters of chloroquine in each cell type.

| Cell type | CC50 | EC50 | TI |
| --- | --- | --- | --- |
| Vero | 134.54 ± 16.76 µM | 9.82 ± 2.79 µM | 13.70 |
| hBMEC | 116.61 ± 9.70 µM | 14.20 ± 0.18 µM | 8.21 |
| hNSC | 94.95 ± 9.38 µM | 12.36 ± 2.76 µM | 7.68 |

CC50 - 50% cytotoxicity concentration; EC50 - half maximal effective concentration; TI - therapeutic index (CC50/EC50). Data are represented as mean ± SD.

A clinical trial of chloroquine administration to DENV-infected patients during three days showed that 60% of the patients in the chloroquine treated group reported feeling less pain and showed improvement in the performance of daily chores during treatment. Symptoms returned after medication withdrawal and chloroquine treatment did not reduce the duration and intensity of the fever or duration of disease (Borges et al., 2013).

Chloroquine is widely distributed to body tissues as well as its analogue hydroxychloroquine. Concentration of hydroxychloroquine in the brain is 4-30 times higher than in the plasma (Titus, 1989). The concentration of chloroquine in the plasma reached 10 μM when a daily intake of 500 mg was prescribed to arthritis patients (Mackenzie, 1983). Chloroquine is able to cross the placental barrier and is supposed to reach similar concentrations on maternal and fetal plasma (Law et al., 2008). Concentrations needed to inhibit ZIKV infection *in vitro*, as shown here, are achieved in the plasma in current chloroquine administration protocols and might even reach the brain. Since it is a molecule already approved for clinical use by the FDA and other agencies around the world, its approval as a therapeutic agent against ZIKV should be faster than new compounds.

The use of chloroquine during pregnancy was evaluated and when prophylactic doses of chloroquine were administered for malaria (400 mg/week), no increment in birth defects was observed (Wolfe and Cordero, 1985). Higher concentrations (250 mg to 500 mg/day) are administered to pregnant women who have severe diseases, such as lupus or rheumatoid arthritis. Few cases of abortion and fetal toxicity were observed. However, fetal toxicity or death could not be discarded as consequence of disease itself since, in most cases, disease was active during pregnancy. In addition, all term deliveries resulted in healthy newborns (Levy et al., 1991; Parke, 1988).

Different mechanisms for chloroquine inhibition of viral infection have been described (Savarino et al., 2003). We observed a higher reduction of ZIKV particles release when the drug was added at 0 hours post-infection, suggesting a higher impact on early stages of infection, possibly during fusion of the envelope protein to the endosome membrane. Chloroquine inhibits acidification of the endosome, consequently inhibiting the low pH-induced conformational changes required for the fusion of the envelope protein of flaviviruses with the endosomal membrane (Smit et al., 2011). However, even when chloroquine was added after early stages of virus infection, except for addition of chloroquine 24 hours post-infection, we noticed a decrease in virus release, suggesting later stages of ZIKV cycle might also be affected, although we cannot eliminate a potential impact on a new cycle of infection.

The proposed mechanism for chloroquine action on later stages of viral replication relates to the alteration of post-translational modifications in the trans-golgi network. Chloroquine treatment impairs *Flavivirus* prM cleavage, which prevents virus maturation and, consequently, infectivity (Randolph et al., 1990; Zybert et al., 2008). For HIV-1, chloroquine inhibits the glycosylation of the gp120 protein, responsible for cell attachment (Tsai et al., 1990). The effect of chloroquine on the glycosylation of ZIKV or other flavivirus envelope protein has yet to be addressed.

ZIKV was detected in the cerebrospinal fluid of ZIKV-infected adult patients that manifested meningoencephalitis, indicating that ZIKV penetrates the central nervous system through yet unknown mechanisms. Transcytosis through the endothelial cells of the blood brain barrier is a known mechanism of viral access to the central nervous system (Dohgu et al., 2012; Suen et al., 2014). Here we demonstrated that chloroquine protects hBMEC, an *in vitro* model of the blood-brain barrier, from ZIKV infection and ZIKV-induced death.

Recent studies showed that neural stem cells are highly permissive for ZIKV infection and one of mechanisms proposed for the cause of microcephaly would be the depletion of the stem cell pool induced by ZIKV (Garcez et al., 2016; Qian et al., 2016; Tang et al., 2016). Our data showed that chloroquine inhibits infection; decreasing the number of induced pluripotent stem cells-derived neural stem cells infected with ZIKV and partially protecting them against ZIKV-induced death. Using the mouse neurospheres model to study neural stem cell differentiation into neurons, another process that might be disturbed in microcephaly, we observed that chloroquine inhibited the infection of neuronal progenitors and partially protected the ability of these cells to extend neurites. The protective effect of chloroquine on stem cells and committed progenitors is potentially a groundbreaking feature of this compound, as it would be prescribed to women at childbearing age traveling to affected countries and women planning pregnancy in endemic areas. This would decrease the chances of infection and thus fetal damage, especially to the developing brain.

Altogether, our results suggest that chloroquine activity against ZIKV should immediately be evaluated *in vivo* and hopefully it will mitigate the devastating brain damage associated with congenital Zika syndrome and neurological damage in affected adults.

## Author contribution

**RD** designed and performed experiments, prepared figures and/or tables, analyzed the data and wrote the manuscript, **LH** designed and performed experiments, prepared figures and/or tables, analyzed the data and wrote the manuscript; **PP** designed and performed experiments, prepared figures and/or tables, analyzed the data and wrote the manuscript; **AV** designed and performed experiments, prepared figures and/ or tables, analyzed the data and wrote the manuscript; **PG** designed and performed experiments, analyzed the data and wrote the manuscript; **FM** designed and performed experiments, prepared figures and/ or tables and analyzed the data; **EL** performed experiments; **SR** designed and performed experiments, contributed with reagents/materials/analysis tools and wrote the manuscript; **LC** designed and performed experiments, prepared figures and/or tables, analyzed the data, contributed with reagents/materials/analysis tools and wrote the manuscript. **RS** designed and performed experiments, contributed with reagents/materials/analysis tools and wrote the manuscript; **AT** designed and performed experiments, prepared figures and/or tables, analyzed the data, contributed with reagents/materials/analysis tools and wrote the manuscript;

## Acknowledgments

This work was supported by Conselho Nacional de Desenvolvimento e Pesquisa (CNPq), Fundação de Amparo à Pesquisa do Estado do Rio de Janeiro (FAPERJ) and Departamento de DST AIDS e Hepatites Virais do Ministério da Saúde do Brasil. The authors acknowledge Manoel Itamar for providing technical support.

**The authors declare no competing financial interests**.

